# Revisiting p53:Sirt1 interaction in the light of controlling p53 acetylation levels

**DOI:** 10.64898/2026.01.19.700258

**Authors:** Vishnupriya Pandey, Frank Hause, Claudio Iacobucci, Christian H. Ihling, Christian Tueting, Panagiotis L. Kastritis, Christian Arlt, Andrea Sinz

## Abstract

The NAD^+^-dependent deacetylase sirtuin 1 (Sirt1) is known to regulate the tumor suppressor p53 via deacetylation, but the structural basis of the protein-protein interaction between full-length p53 and Sirt1 has so far remained elusive. We apply an integrated approach, combining structural mass spectrometry (MS) with data-driven molecular docking, to study the interaction between human p53 and Sirt1. Sirt1 was found to bind exclusively to acetylated p53 forming complexes with a 1:1 stoichiometry, irrespectively of p53’s oligomeric state. The lysine residue at position 582 (K582) in p53 was identified as predominant acetylation site showing a selective Sirt1-dependent deacetylation at this position. Cross-linking mass spectrometry (XL-MS) provided valuable distance constraints between p53 and Sirt1. Specifically, cross-links created between p53-K382 / Sirt1-K427 and p53-K120 / Sirt1-K622 give hints on a highly flexible interface. Molecular docking was conducted based on the distance constraints imposed by the cross-links, positioning Sirt1 at the DNA-binding and tetramerization domains of p53. This gives a rationale for a steric exclusion of additional Sirt1 molecules binding to p53. We present the first structural model of the full-length p53:Sirt1 (1:1) complex, establishing a mechanistic framework that links p53 activity to its Sirt1-controlled acetylation status.

## Introduction

The homotetrameric tumor suppressor p53 is a transcription factor orchestrating crucial cellular processes, including cell cycle arrest, DNA repair, apoptosis, and senescence (ref. 1, 2). p53 possesses extensive regions of intrinsic disorder, categorizing it as an intrinsically disordered protein (IDP) (ref. 3, 4). The functional diversity of p53 is regulated by post-translational modifications (PTMs), with acetylation emerging as a particularly significant modification. Acetylation, predominantly mediated by histone acetyltransferases (HATs), such as p300, occurs at distinct lysine residues within p53 and substantially influences p53’s stability, DNA-binding specificity, transcriptional activity, and interactions with regulatory cofactors (ref. 5–7). This modification is dynamically reversed by the NAD⁺-dependent deacetylase sirtuin 1 (Sirt1), which removes acetyl groups in a site-specific manner, thereby establishing a balance between acetylation and deacetylation that governs the structural and functional states of p53 (ref. 8–10).

The aim of this study is to elucidate the conformational and stoichiometric characteristics underlying the acetylation-dependent p53:Sirt1 interaction. We specifically examine whether the interaction between acetylated p53 variants and Sirt1 occurs independently of p53’s oligomeric states using monomeric (p53_L344P_), dimeric (p53_L344A_), and tetrameric (p53_wt_) forms (ref. 11).

To comprehensively address this question, we purified recombinant protein variants of p53 and Sirt1 and studied them by state-of-the-art techniques of structural mass spectrometry (MS). As such, native MS represents a robust analytical technique for studying protein complexes under near native conditions by preserving non-covalent interactions. This allows analyzing intact protein assemblies to infer their subunit compositions and stoichiometries (ref. 12–14). p53 was subjected to targeted enzymatic acetylation by p300 before applying native MS, XL-MS, and computational modeling approaches. Leveraging native MS, we systematically investigate interactions between acetylated p53 variants (^Ac^p53) and Sirt1, elucidating how acetylation patterns and oligomeric states of p53 influence the formation and stability of these protein complexes. Additionally, chemical cross-linking coupled with mass spectrometry (XL-MS) (ref. 15–17) and molecular docking provide complementary structural insights, allowing a precise identification of interacting residues and mapping the interfaces between ^Ac^p53 variants and Sirt1.

Our integrative approach facilitates the identification of critical molecular determinants that govern p53:Sirt1 interaction. We provide valuable insights into the biological significance of this protein-protein interaction shedding light on its influence regarding cell fate decisions.

## Materials and Methods

### Expression and purification of human Sirt1 and human p53 variants

Human sirtuin 1 (Sirt1) was expressed in *E. coli* BL21(DE3) cells as N-terminally truncated version (SIRT1_Δ6–83_) (ref. 18). The codon-optimized sequence was synthesized (Invitrogen) with N-terminal 6×His tag and thrombin cleavage site and cloned into a pET-28a(+) vector. Cells were grown to OD_600_ 0.7–0.8, induction was performed with 1 mM IPTG, and cells were incubated at 25 °C for 6 h before they were pelleted (6,000×g, 12 min, 4 °C) and stored at −20 °C.

Pellets were resuspended in lysis buffer (50 mM HEPES, 250 mM NaCl, 5 mM imidazole, 1 mM TCEP, pH 8.0) with protease inhibitors, lysed by ultrasonication, treated with benzonase (2.5 U/mL) and 3 mM MgCl_2_, and incubated on ice for 30 min. Clarified lysate (25 000×g, 30 min, 4 °C) was subjected to immobilized metal-ion affinity chromatography (1-mL HisTrap HP column, Cytiva) with a 5–200 mM imidazole gradient on an ÄKTA Pure system (Cytiva). Fractions were pooled and subjected to size-exclusion chromatography (HiLoad Superdex 200 pg, 16/600 column, Cytiva) in 25 mM HEPES, 100 mM NaCl, 2.5 mM TCEP, 10% glycerol, pH 7.5. Purified Sirt1 was concentrated to 15 µM (Amicon Ultra 50 kDa MWCO) and stored at −80 °C. Sirt1 activity was monitored with the Universal SIRT1 Activity Assay Kit (Abcam).

Monomeric (p53_L344P_) and dimeric (p53_L344A_) p53 variants were expressed and purified following an established protocol (ref. 19). For acetylated p53_wt_ (^Ac^p53_wt_), the p300 HAT domain (residues 1284–1673) was codon-optimized for *E. coli*, cloned into a pBAD30 vector, and co-expressed with p53 in BL21(DE3). Induction was performed with 0.5 mM IPTG and 0.1% L-arabinose.

### Cross-linking reactions and enzymatic digestion

Cross-linking was performed with p53 and Sirt1 (5 µM each) in 10 µM NAD^+^ in 50 mM HEPES, 300 mM NaCl, 2.5 mM TCEP, 10% glycerol, pH 7.2. The cross-linker DSBU (disuccinimidyl dibutyric urea) (ref. 20) was dissolved in DMSO directly before the cross-linking reaction and added at 50-fold molar excess over total protein concentration. Cross-linking reactions were performed at 4°C and quenched after 60 min with 20 mM ammonium bicarbonate. In-gel and in-solution enzymatic digestions were performed as described previously (ref. 21).

### LC-MS/MS of cross-linked peptides

Fractionation of proteolytic peptide mixtures was carried out on an UltiMate 3000 RSLC nano-HPLC system (Thermo Fisher Scientific) using a reversed-phase C18 precolumn (Acclaim PepMap 100, 300 μm ID × 5 mm, 5 μm, 100 Å) and a Picofrit C18 separation column (75 µm ID x 40 cm, Tip ID 15 µm, New Objective, packed with Reprosil-Pur 120 C18-AQ, 3 µm, 120 Å, Dr. Maisch GmbH). After washing the peptide mixture on the precolumn (water containing 0.1% (v/v) TFA) for 15 min, peptides were eluted onto the separation column and eluted with a linear gradient of 3% to 50% solvent B in solvent A (120 min), 50% to 85% B (5 min), and 85% B (5 min); solvent A: water containing 0.1% formic acid and solvent B: acetonitrile containing 0.1% formic acid. The nano-HPLC column was directly coupled to a CaptiveSpray source of timsTOF Pro mass spectrometer (Bruker Daltonik). Data were acquired in data-dependent MS/MS mode and peptides were analyzed with a parallel accumulation serial fragmentation (PASEF) method for standard proteomics. Precursor ions (*m/z* 100–1700, charge states 3+ to 8+) were selected for trapped ion mobility spectrometry (TIMS) separation and fragmentation (scan range 0.6–1.6 Vs/cm^2^, collision energy ramping between 95 eV at 1.6 Vs/cm^2^ (1/k0 End) and 20 eV at 0.6 Vs/cm^2^ (1/k0 Start), ramp time 200 ms, ramp rate 4.85 Hz, accumulation time 200 ms, duty cycle 100 %, 10 PASEF ramps per cycle (2.47 s), target intensity per PASEF precursor 100,000 with intensity threshold 1,000, active exclusion enabled).

### Cross-link analysis

For the identification of cross-links, MS/MS data were analyzed using MeroX (version 2.0.1.7) (ref. 22). An FDR cut off of 1% and a MeroX cut-off score of 40 were applied. All identified intra-protein, inter-protein, and “dead-end” cross-links were manually validated and visualized via Circos plots (Circos, (ref. 23)). XL-MS data are summarized in Supplementary Table I.

### Native MS of intact proteins

Native MS experiments were carried out on a modified high-mass QToF II instrument (Waters Micromass/MS Vision) using p53 and Sirt1 solutions of 10 µM in 500 mM ammonium acetate, pH 6. For p53:DNA complexes, ^Ac^p53_wt_ was mixed (1:1.2) with double-stranded DNA response element (RE: CGCGGACATGTCCGGACATGTCCCGC) (ref. 24) and incubated at 4 °C overnight; for p53:Sirt1 complexes, ^Ac^p53 and Sirt1 were 1:1 mixed and incubated at 4 °C for 1 h before native MS experiments were conducted.

The following MS settings were applied: capillary voltage 1300–1400 V, cone voltage 130–160 V, extraction voltage 15–30 V, collision energies 60–80 V, source pressure 1.0 x 10^-1^ mbar, collision cell pressure 1.0–1.3 x 10^-2^ mbar. The instrument was operated in manual MS profile mode at an *m/z* range of 1,000–10,000. For collision-induced dissociation tandem mass spectrometry (CID-MS/MS) experiments, signals at specific *m/z* values were isolated and fragmented by applying a collision energy of 120-150 V.

### p53 acetylation status

To determine p53’s acetylation status, purified ^Ac^p53 (2 µg) was denatured by adding 8 M urea in 400 mM ammonium bicarbonate. ^Ac^p53 was reduced with dithiothreitol (DTT) and alkylated with iodoacetamide (IAA). To determine deacetylation rates of ^Ac^p53 by Sirt1, equimolar concentrations of ^Ac^p53 and Sirt1 of 5 µM each were incubated for 0, 2, 5, 10, 20 and 40 min in 8 M urea, 400 mM ammonium bicarbonate in presence of 10 µM NAD^+^. LC-MS/MS analysis was conducted after enzymatic digestion of the proteins. Proteinase Arg-C (sequencing grade, Promega) was added at an enzyme-to-substrate ratio of 1:55 (w/w), and digestion was carried out at 37 °C overnight. Arg-C digested peptides were separated with an UltiMate 3000 RSLC nano-HPLC system (Thermo Fisher Scientific) and analyzed using an Orbitrap Fusion Tribrid mass spectrometer with EASY Spray source (Thermo Fisher Scientific) according to an existing protocol (ref. 4). For quantifying acetylation sites in p53, extracted ion chromatograms were generated for all modification states of peptides containing K370, K372, K373, K381, K382, and K386. Charge states between 2+ to 5+ were considered and abundances were added up. To determine relative acetylation levels at each lysine residue, the abundances of p53 peptides were normalized to all versions of the respective p53 peptides that were detected.

### Molecular Docking

For molecular docking, the HADDOCK 2.4 webserver was used (ref. 25). The input models were derived from the AlphaFold2 database (ref. 26) and models are accessible under the accession codes “AF-P04637-F1-model_v4” for p53 and “AF-Q96EB6-F1-model_v4” for Sirt1. Prior to rigid-body docking, low-confidence regions, based on plDDT score and domain proximity, were removed. For docking, the following parts of the models were used: Sirt1183-670;Δ501-633, p53DBD;95-295 and p53TET;320-360. Cross-links were defined as constraints between ClZ atoms at a distance range of 5–25 Å. Cross-links between the tetramerization domain and residue 633 in Sirt1, which is not included in the docked model due to a low prediction confidence, were mapped to residue 621 of Sirt1 at a distance range of 5–30 Å, accounting for the high flexibility of the unresolved region 621–632. Primary sequence constraints were defined between residue 274 of p53’s DNA binding domain (DBD) and residue 320 of p53’s tetramerization domain (TET), and between residue 360 of the p53 tetramerization domain and the active site of Sirt1 at residue 363 (where K382 gets deacylated) at a distance range of 0–30 Å. All distance constraints are listed in Supplementary Table II. All constraints were used as unambiguous constraints and docking was performed in multi-body docking (ref. 27) by default. All 200 water-refined structures were aligned to Sirt1 and 12 outlier structures were removed manually. Alignment and visualization were done in ChimeraX (ref. 28) and PyMOL (ref. 29).

### Data deposition

All MS data generated in this study have been deposited to the ProteomeXchange Consortium via the PRIDE partner repository with the project accession PXD072665, username, password: 0rEy6XiNVJXk.

## Results

### Expression of p53 and ^Ac^p53 variants and Sirt1

p53 and ^Ac^p53 variants were expressed and purified according to an existing protocol (ref. 19) and p53 acetylation was investigated by native MS. The native mass spectrum of ^Ac^p53_L344P_ revealed different acetylation states of the monomer with a small degree of dimer (Fig. 1A). In contrast, the native mass spectrum of ^Ac^p53_L344A_ showed a prominent dimer with acetylations at seven residues, whereas monomeric p53_L344A_ displayed a mixture of non-acetylated and singly or doubly acetylated forms (Fig. 1B). For ^Ac^p53_wt_, the tetramer with six acetylated lysine residues was observed among other oligomeric species (Fig. 1C). The *E. coli* chaperone DnaK was consistently observed in all spectra of p53_wt_ (Figs. 1C, S1, S2).

Notably, both p53_wt_ and ^Ac^p53_wt_ were capable of DNA binding, as evidenced by their ability to bind to a DNA response element (RE, 26 bp), with the ^Ac^p53_wt_:DNA complex being detected in the native mass spectrum (Fig. S2). ^Ac^p53_wt_ was predominantly detected as a tetramer bound to DNA RE (Fig. S2B) in contrast to p53_wt_, which was observed as monomeric, dimeric, and tetrameric species (Fig. S2A).

Sirt1 was expressed in *E. coli* and its identity was confirmed by peptide fragmentation analysis via LC-MS/MS. Native MS shows the presence of intact Sirt1 in monomeric (charge states 15+ to 19+) and dimeric (charge states 23+ to 28+) forms (Fig. S3).

### The impact of p53’s acetylation and oligomeric states on p53:Sirt1 interaction

To assess how acetylation and oligomeric states influence complex formation, monomeric (p53_L344P_), dimeric (p53_L344A_), and tetrameric (p53_wt_) p53 variants were incubated with Sirt1 in both acetylated and non-acetylated forms and analyzed by native MS. Complexes were consistently observed only with ^Ac^p53, independent of its oligomeric state, while non-acetylated variants did not interact with Sirt1. The detected masses of the complexes indicate a consistent one-to-one stoichiometry, with a single Sirt1 molecule binding to either an ^Ac^p53 monomer (Fig. S4), dimer (Fig. S5), or tetramer (Fig. 2).

The formation of ^Ac^p53:Sirt1 complexes was further confirmed by collisional activation (CID-MS/MS, Figs. 2C, S4C, S5C). Interestingly, dissociation of all complexes uniformly yielded ^Ac^p53 monomers and ^Ac^p53:Sirt1 species containing one, two, or three ^Ac^p53 molecules bound to one Sirt1 molecule.

### Site-specific acetylation and deacetylation of p53

In-depth proteome analyses of p53 were employed to characterize the lysine residues that were acetylated by p300. LC-MS/MS analyses of ^Ac^p53_wt_ confirmed acetylation at lysine residues K370, K372, K373, K381, K382, and K386 (Fig. 3). As single acetylations at K370, K372, and K373 could not be unambiguously pinpointed to one specific residue, these three lysine residues were collectively analyzed and mono-, di-, and tri-acetylated species were quantified as combined K370/K372/K373 states. Acetylation was observed to be most prominent for the mono-acetylated state (67–86%) at K382 (51–84%), while K381 showed moderate acetylation (15–19%) and K386 remained mainly unmodified (<2%). These acetylation patterns are in good agreement with previous findings in cell culture (human lung carcinoma cells H460) where the same lysines were found to be acetylated by p300 (ref. 30).

To examine the site-specificity of Sirt1 activity, we monitored deacetylation kinetics for lysine residues K381 and K382. For all three variants, monomeric p53_L344P_, dimeric p53_L344A_, and tetrameric p53_wt_, a time-dependent decrease in mono-acetylation at K382 was observed, accompanied by a corresponding increase in the signal of unmodified p53. Our data show a minor decrease of mono-acetylation at K381 and a site-specific deacetylation of K382 by Sirt1 in a time-dependent manner (Fig. 4).

### Molecular details of p53:Sirt1 interaction

To characterize the molecular interface between ^Ac^p53 and Sirt1 we performed XL-MS (ref. 15) that covalently connects interacting regions of proteins. Acetylated and non-acetylated p53 variants were chemically cross-linked with Sirt1 using the homobifunctional, amine-reactive cross-linker DSBU (ref. 20). SDS-PAGE of the cross-linked products revealed that Sirt1 forms complexes only with ^Ac^p53, consistent with native MS findings (Fig. 5). Specifically, cross-linking of ^Ac^p53_L344P_ with Sirt1 produced a complex at an apparent molecular weight of ∼220 kDa, representing a single Sirt1 molecule interacting with an ^Ac^p53_L344P_ monomer. In contrast, no interaction was observed when Sirt1 was incubated with non-acetylated p53_L344P_ (Fig. 5A).

Similarly, cross-linking of Sirt1 with dimeric ^Ac^p53_L344A_ resulted in a complex at an apparent molecular weight of ∼250 kDa, corresponding to a single Sirt1 molecule cross-linked to an ^Ac^p53_L344A_ dimer. No complex was formed with non-acetylated p53_L344A_ (Fig. 5B). In the case of tetrameric ^Ac^p53_wt_, the cross-linking reaction produced a higher-molecular weight species, potentially corresponding to a single Sirt1 molecule cross-linked to an ^Ac^p53_wt_ tetramer. Again, no interaction was observed for non-acetylated p53_wt_ (Fig. 5C).

XL-MS analyses identified ten unique inter-protein cross-linking sites for ^Ac^p53_L344P_, five for ^Ac^p53_L344A_, and seven for ^Ac^p53_wt_ (Fig. 6, Supplementary Table I).

An inter-protein cross-link that was found for all three p53 variants involves K382 in the C-terminal regulatory region of p53 and K427 in the catalytic domain of Sirt1. A second recurrent cross-link is p53-K120:Sirt1-K622. As K120 is not acetylated by p300, but rather by Tip60/MOF in the cell (ref. 31–33), it remains unmodified in our experimental setup and is therefore available for cross-linking. Notably, all cross-links identified involved at least one residue located in an intrinsically disordered region (IDR).

The XL-MS results provide detailed molecular insights into the p53:Sirt1 interaction: As p53-K382 cannot form a cross-link as long as it remains acetylated, the p53-K382:Sirt1-K427 cross-link suggests that it is created immediately after removal of the acetyl group when the two proteins are still in close spatial proximity. This is consistent with the deacetylation kinetics indicating the site-specific removal of the acetylation at p53-K382 by Sirt1, and explains why the K382-K427 cross-link was detected across all oligomeric states of p53.

Together, these data suggest that acetylation at K382 is a mandatory prerequisite for initial p53:Sirt1 complex formation, while residues, such as K120, provide additional stabilizing contacts independent of the acetylation status.

The distance constraints imposed by the cross-links (Supplementary Table II) were then subjected to computational modeling to deduce valuable insights into a full-length 3D-structural model of the p53:Sirt1 (1:1) complex.

### Computational modeling to derive 3D-structural data of full-length p53:Sirt1 complex

For computational modeling, a rigid-body docking approach was employed using HADDOCK (ref. 25) and the distance constraints derived from XL-MS. AlphaFold2-predicted monomeric structures of Sirt1, the DNA-binding domain of p53 (p53_DBD_), and the tetramerization domain of p53 (p53_TET_) served to generate a 3D-structural model of the 1:1 complex. Low-confidence regions in the AlphaFold2 predictions were removed prior to docking and additional distance constraints were derived from the X-ray structure of an acetylated C-terminal p53 peptide bound to Sirt1 (ref. 34) (Fig. S6). Clustering analysis retained a total of 188 out of 200 water-refined models and their median pairwise backbone root mean square deviation (RMSD) was 3.7 ± 1.3 Å (SD), indicating high confidence in the data-driven docking results. Specifically, p53_DBD_ and p53_TET_ clustered at distinct regions of Sirt1 (Fig. 7A). Electrostatic interactions, mediated by polar and charged residues, dominate the p53:Sirt1 interface (Fig. 7B-D). The models derived exhibit a high satisfaction of the experimentally measured cross-links (Fig. 7E), highlighting the critical roles of p53_DBD_ and p53_TET_ in binding and coordinating Sirt1.

## Discussion

Despite extensive research on the regulatory interplay between p53 and Sirt1, structural and mechanistic details of their interaction have remained elusive to date. Full-length p53 comprises large IDRs accounting for circa 57% (ref. 35) of the total protein, predominantly at its N- and C-termini. Sirt1 exhibits comparable structural plasticity, with also circa 57% (ref. 35) of its sequence predicted to be intrinsically disordered, particularly in the N-terminal regulatory region and the inter-domain linkers. This high degree of disorder renders both proteins challenging to high-resolution analysis by X-ray crystallography while size-dependent, they are also challenging for current cryo-EM analysis. Consequently, previous structural insights have been confined to short C-terminal p53 peptides (residues 382–396) in complex with the Sirt1 catalytic domain (ref. 34). To overcome these limitations and achieve a physiologically more relevant view of the interaction, we studied full-length human p53 and Sirt1 across defined p53 oligomeric states via an integrated structural MS workflow.

Taken together, our results indicate that Sirt1 associates with p53 exclusively in its acetylated form, forming a complex with an apparent 1:1 stoichiometry per p53 molecule, independent of p53’s oligomeric state. Acetylation at K382 emerged as the principal determinant for Sirt1 recruitment, complemented by additional stabilizing interactions mediated through IDRs of both proteins. Notably, our study provides a three-dimensional structural model of the full-length p53:Sirt1 (1:1) complex where polar interactions dominate the interface.

### p53:Sirt1 interaction is independent of p53 stoichiometry

Native MS consistently detected a single Sirt1 molecule bound to ^Ac^p53 across monomeric, dimeric, and tetrameric states, while non-acetylated p53 did not engage in Sirt1 interaction (Fig. 2, S4, S5). Dissociation experiments yielded ^Ac^p53 monomers and ^Ac^p53:Sirt1 subcomplexes, supporting the presence of a defined binding interface rather than a diffuse surface association. The oligomerization-independent 1:1 stoichiometry indicates that a single Sirt1 molecule per p53 molecule is sufficient to initiate deacetylation, suggesting that steric constraints or an induced-fit rearrangement limit the simultaneous binding of additional Sirt1 molecules. This interpretation is consistent with our computational modeling results showing that Sirt1 engagement restricts the relative mobility of the p53_DBD_ and p53_TET_ domains, thereby confining their spatial organization within the complex (Fig. 7). Such steric and conformational restrictions hint a sequential, processive mode of deacetylation, in which one Sirt1 molecule acts on an individual p53 monomer before dissociating and re-engaging another.

Strikingly, an analogous oligomerization-independent stoichiometry has been described for S100β where one S100β dimer associates with each p53 molecule, irrespective of its oligomeric state (ref. 11). However, in contrast to Sirt1, S100β binding can destabilize p53 oligomers, shifting the equilibrium toward monomeric p53 and thereby attenuating its transcriptional activity (ref. 36). These parallels suggest that oligomerization-independent binding might represent a broader regulatory mechanism, by which partner proteins fine-tune p53 activity through conformational modulation rather than direct competition for binding sites.

### K382 acts as principal acetylation site and functional control point

K382 is the predominant acetylation site of p53 across all oligomeric states (Fig. 3). In the cellular environment, p53 predominantly exists as dimer under basal conditions and assembles into a tetramer only upon stress or DNA damage (ref. 37, 38). Acetylation at K382 might already occur in lower-order oligomers, potentially priming p53 for activation and tetramerization.

Acetylation occurs with high efficiency, reaching ca. 70% in monomeric and dimeric p53 and 92% in the tetrameric form (Fig. 3). Time-resolved deacetylation kinetics further revealed that co-incubation of acetylated p53 with Sirt1 led to a rapid and selective loss of the acetyl group at residue K382, while deacetylation at the adjacent residue K381 was comparatively modest (Fig. 4). These results identify K382 as the primary target for Sirt1 and underline its role as central regulatory site governing p53 deacetylation. K382 has been recognized as a major regulatory hotspot within the p53 C-terminus, serving as a convergence point for multiple PTMs, including methylation (ref. 39), acetylation (ref. 5), and ubiquitination (ref. 40). Previous studies have shown that acetylation at K382 enhances DNA binding (ref. 5), transcriptional activation (ref. 41), and protein stability (ref. 42), while its modification state dictates p53’s functional outcome in stress signaling (ref. 43). Our data corroborate and extend these earlier findings by demonstrating that K382 acetylation occurs efficiently across all p53 oligomeric states and represents the principal site of Sirt1-catalyzed deacetylation in the context of the full-length p53:Sirt1 complex.

Our observations differ from those of Itahana *et al.* (ref. 44) who reported inefficient acetylation of monomeric and dimeric p53 in mammalian cells. This discrepancy likely arises from the different experimental systems: p53 expressed in *E. coli* lacks mammalian cofactors that might restrict acetylation of lower oligomeric forms. The multiple mutations introduced in that previous study (ref. 44) within the nuclear export and tetramerization domains (L348A/L350A/M340A/L344A) apparently interfere with C-terminal acetylation. In addition, an altered subcellular localization of the quadruple mutant in a mammalian context may further hinder access of nuclear acetyltransferases, an effect absent in the prokaryotic expression system used here.

Collectively, our findings confirm K382 as a pivotal post-translational switch integrating acetylation dynamics with p53’s structural and functional plasticity.

### Integrative structural analysis of the p53:Sirt1 complex

Consistent with the site-specific deacetylation kinetics, XL-MS identified a recurrent inter-protein cross-linking site between p53-K382 and Sirt1-K427 that was detected across all p53 oligomeric states. This cross-link connects the C-terminal regulatory region of p53 with the catalytic domain of Sirt1 and indicates direct spatial proximity during catalysis. A second recurrent constraint links p53-K120 in the DNA-binding domain to Sirt1-K622. As K120 is acetylated in mammalian cells by Tip60/MOF rather than by p300 (ref. 31–33), it remains unmodified in our experimental system and is therefore accessible for cross-link formation. Notably, all inter-protein cross-links identified involved at least one residue located within an IDR, highlighting the flexible and adaptive nature of the p53:Sirt1 interface.

The XL-MS results provide molecular insights into a potential interaction mechanism between p53 and Sirt1 (Fig. 6). As acetylated K382 of p53 cannot participate in cross-link formation, the detection of the p53-K382:Sirt1-K427 linkage implies that it is formed immediately after deacetylation when both proteins are in close spatial proximity. This observation aligns with the rapid, site-specific loss of acetylation at K382 observed in kinetic assays and explains the consistent presence of the p53-K382:Sirt1-K427 cross-link across all oligomeric states. Together, these data indicate that acetylation at K382 is a prerequisite for initial p53:Sirt1 (1:1) complex formation, as confirmed by native MS. Residues, such as K120, provide additional stabilizing contacts that are independent of acetylation status. The complete XL-MS dataset (Supplementary Table I) yielded distance constraints for computational modeling to derive a structural model of the full-length p53:Sirt1 (1:1) complex.

Both p53 and Sirt1 contain a large degree of IDRs, allowing them to adopt flexible conformational ensembles in solution. Consequently, the binding of the 77-kDa Sirt1 monomer to the 87-kDa p53 dimer, or the 175-kDa p53 tetramer, is likely to impose steric restrictions that preclude the simultaneous binding of additional Sirt1 molecules to the same complex. In this setting, a single Sirt1 molecule deacetylates one p53 monomer and then dissociates, after which it can rebind to and deacetylate additional monomers within the same p53 tetramer. Alternatively, one Sirt1 molecule could undergo local repositioning within the complex to catalyze deacetylation on adjacent p53 monomers without full dissociation.

Docking simulations suggest that monomeric Sirt1 engages both with the p53_DBD_ and p53_TET_ domains simultaneously (Fig. 7), constraining their relative spatial orientations. In the unbound state, the p53_DBD_ and p53_TET_ domains are connected via a flexible linker, allowing substantial conformational freedom. While additional binding sites within higher-order p53 assemblies will remain individually accessible, the combined interface accommodating one Sirt1 molecule becomes sterically saturated. When the rigid-body-docked p53:Sirt1 (1:1) complex is superimposed on the p53_TET_ crystal structure (PDB 1C26 (ref. 45)), a theoretical 4:4 stoichiometry is geometrically possible, but this would result in distances between p53_DBD_ molecules of 84 to 114 Å, separated by a central planar p53_TET_:Sirt1core (Fig. S7). Such an arrangement is inconsistent with known p53 conformations, supporting a model, in which binding of a single Sirt1 molecule sterically and conformationally precludes a stable association of additional Sirt1 units.

Together, the XL-MS constraints and docking results reveal a dynamic 1:1 interaction, in which Sirt1 interacts with acetylated p53 through an interplay of an ordered interface and flexible, IDR-mediated contacts and thereby regulates sequential deacetylation cycles within the p53 complex.

## Acknowledgements

This work was supported by the Deutsche Forschungsgemeinschaft (DFG) (RTG 2467, project number 391498659 “Intrinsically Disordered Proteins-Molecular Principles, Cellular Functions, and Diseases”, CRC 1664, project number 514901783 “SNP2Prot - Plant Proteoform Diversity”, and CRC 1423, project number 421152132 “Structural Dynamics of GPCR Activation and Signaling”). AS acknowledges additional financial support by the DFG (RTG 2751 “InCuPanC”, project number 449501615; RU 5433 “RNA in Focus” project number 468534282, INST 271/404-1 FUGG; INST 271/405-1 FUGG; INST271/528-1 FUGG), the Federal Ministry for Economic Affairs and Energy (BMWi, ZIM project KK5096401SK0), the region of Saxony-Anhalt, and the Martin Luther University Halle-Wittenberg (Center for Structural Mass Spectrometry). PLK acknowledges funding by the European Union through funding of the Horizon Europe ERA Chair “hot4cryo” project number 101086665, the Federal Ministry for Education and Research (BMBF, ZIK program; grant numbers 03Z22HN23 and 03COV04), and the European Regional Development Funds for Saxony-Anhalt (grant numbers EFRE: ZS/2016/04/78115 and ZS/2024/05/187255).

## Author Contributions

VP performed the experimental procedures, analyzed the data, generated visualizations, and contributed to manuscript writing. FH analyzed the data, generated visualizations, participated in the conceptualization of the work, wrote and reviewed the manuscript. CI performed experimental procedures and analyzed MS data. CHI and CA performed MS and XL-MS data analysis. CA participated in the conceptualization of the work. CT conducted molecular modeling and contributed to visualization. PLK contributed to molecular modeling, reviewed the manuscript, and acquired funding. AS conceptionalized this work, wrote, and reviewed the manuscript, and acquired funding.

## Data Availability Statement

The datasets generated during and/or analysed during the current study are available in the ProteomeXchange Consortium via the PRIDE partner repository with identifier PXD072665.

---

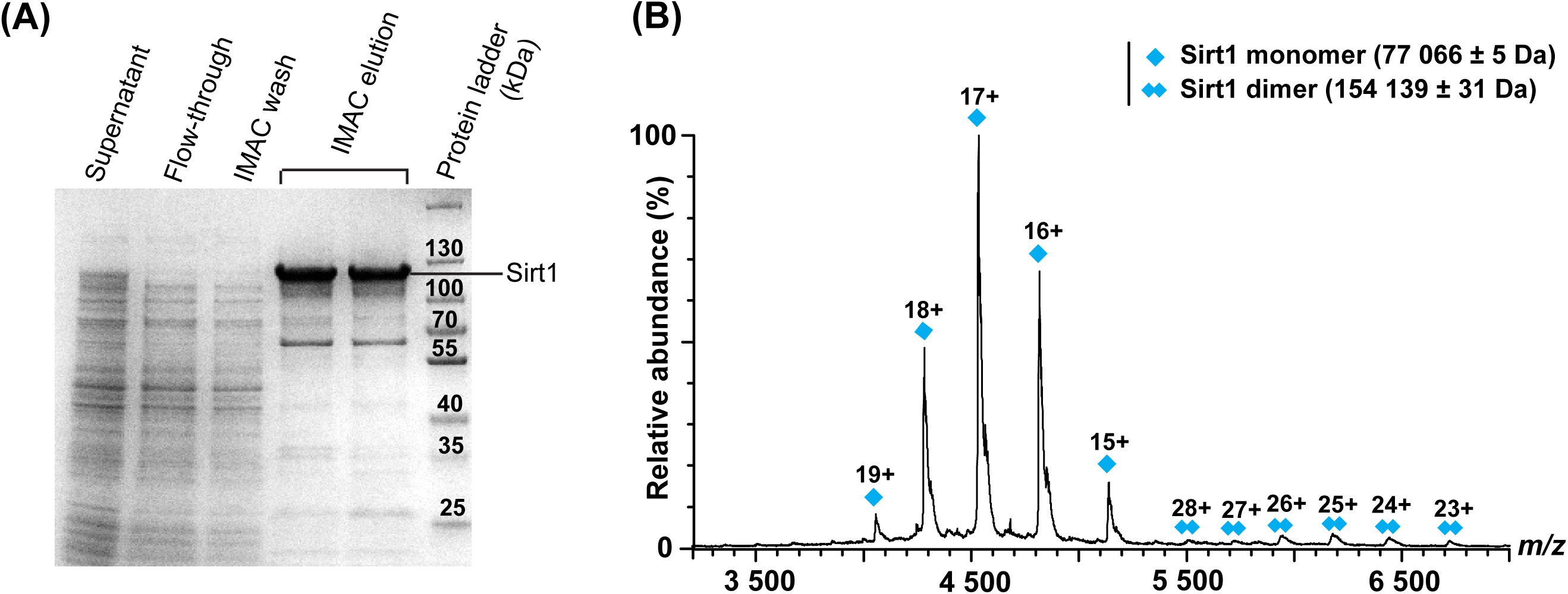

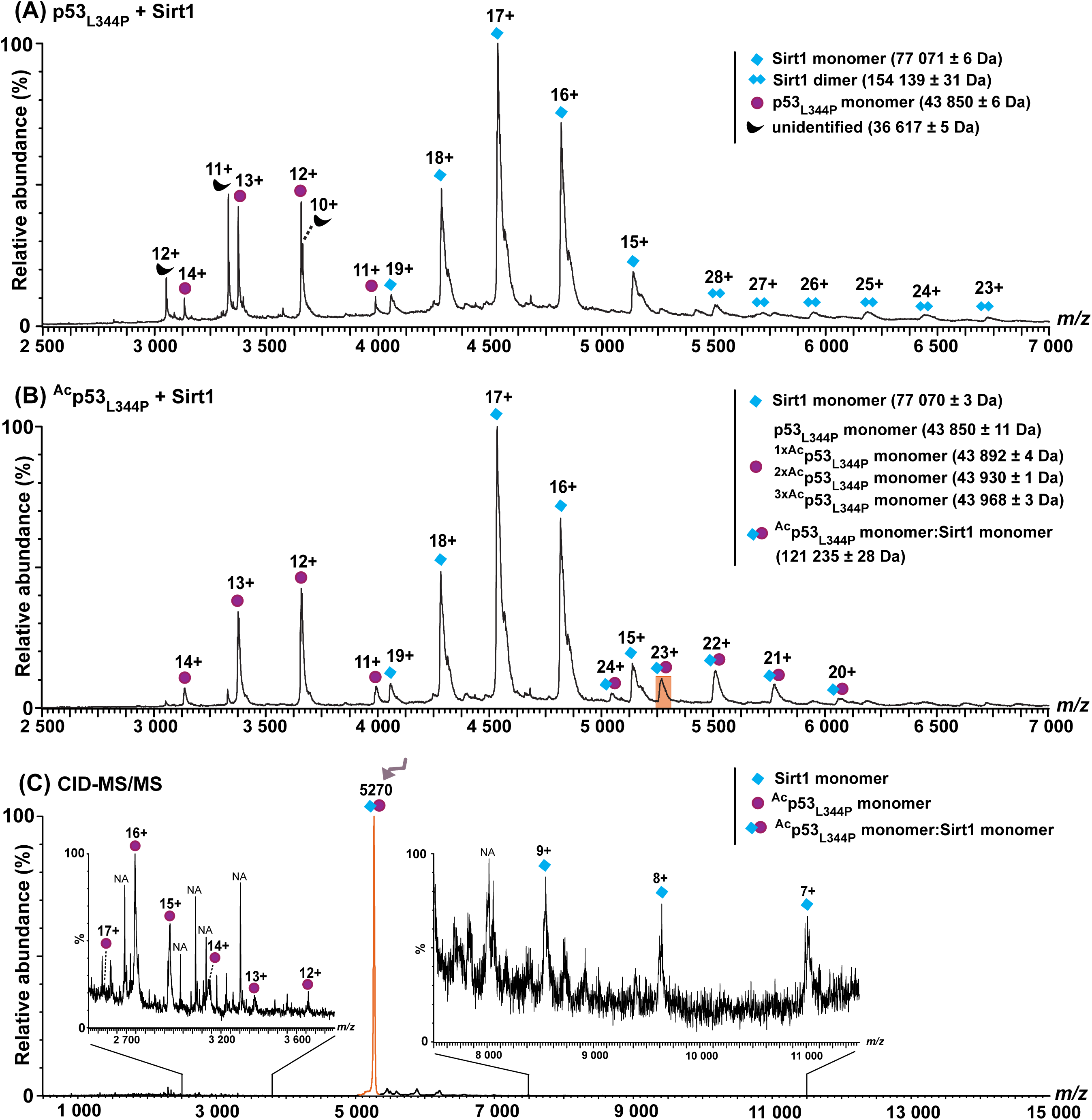

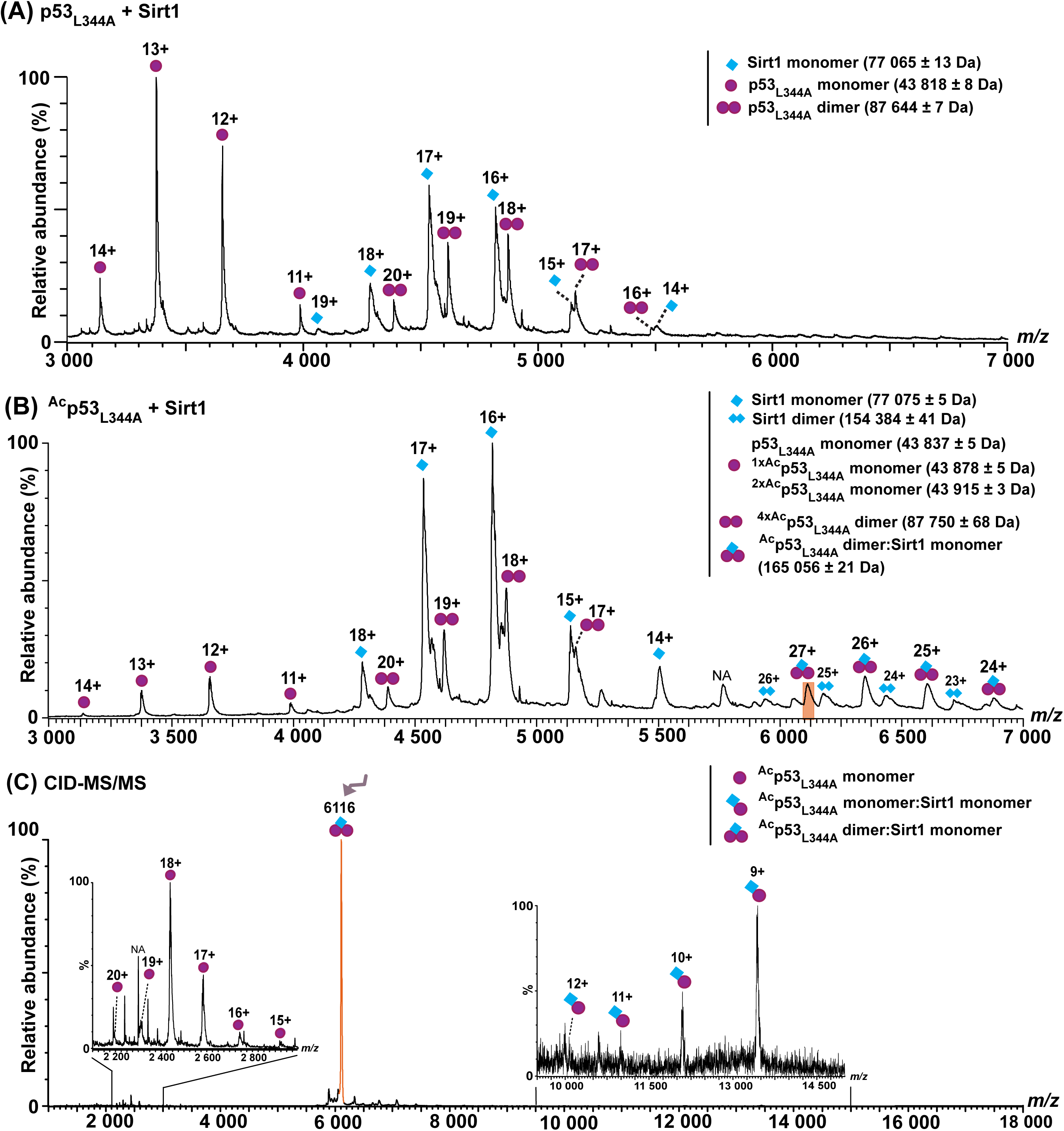

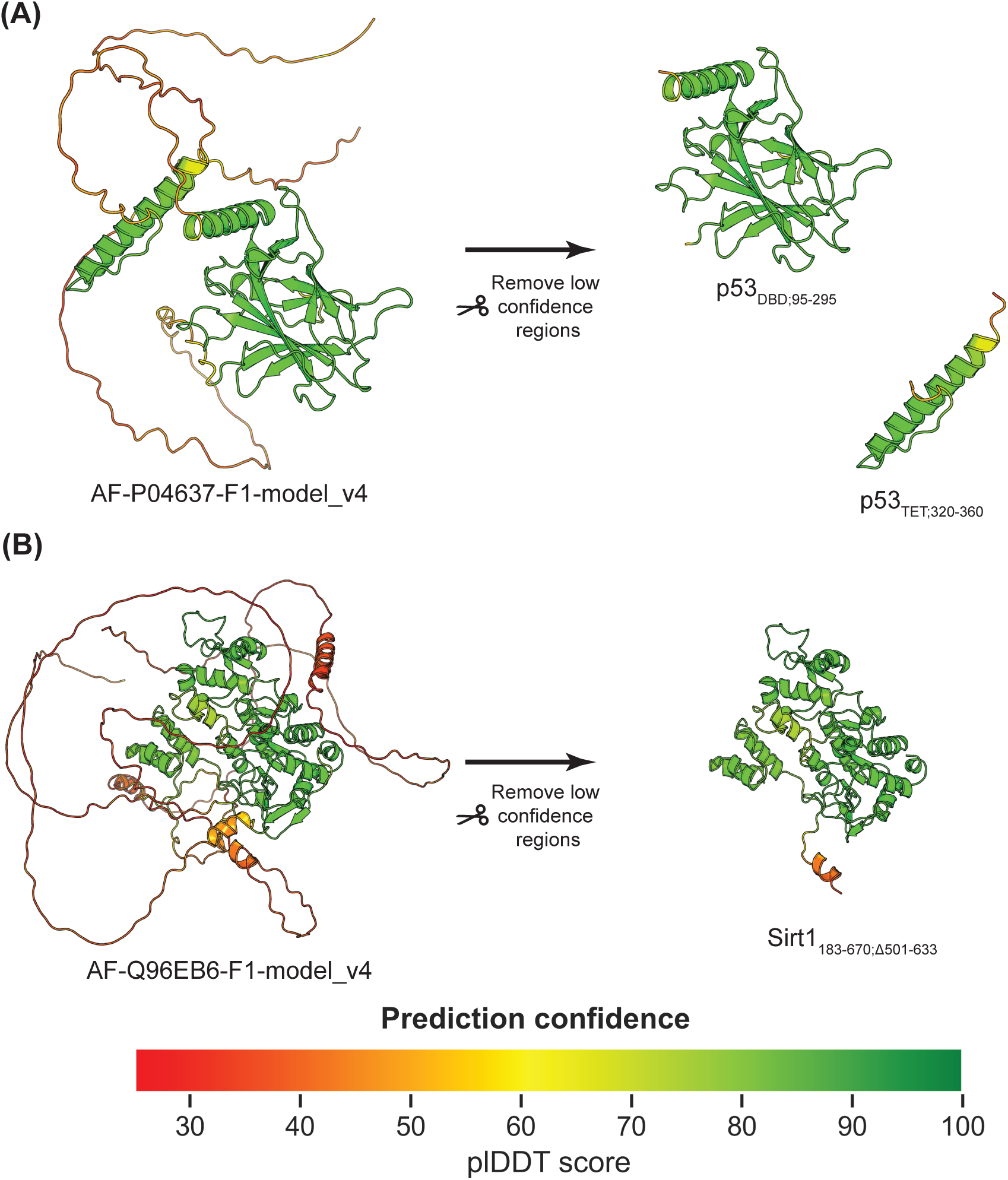

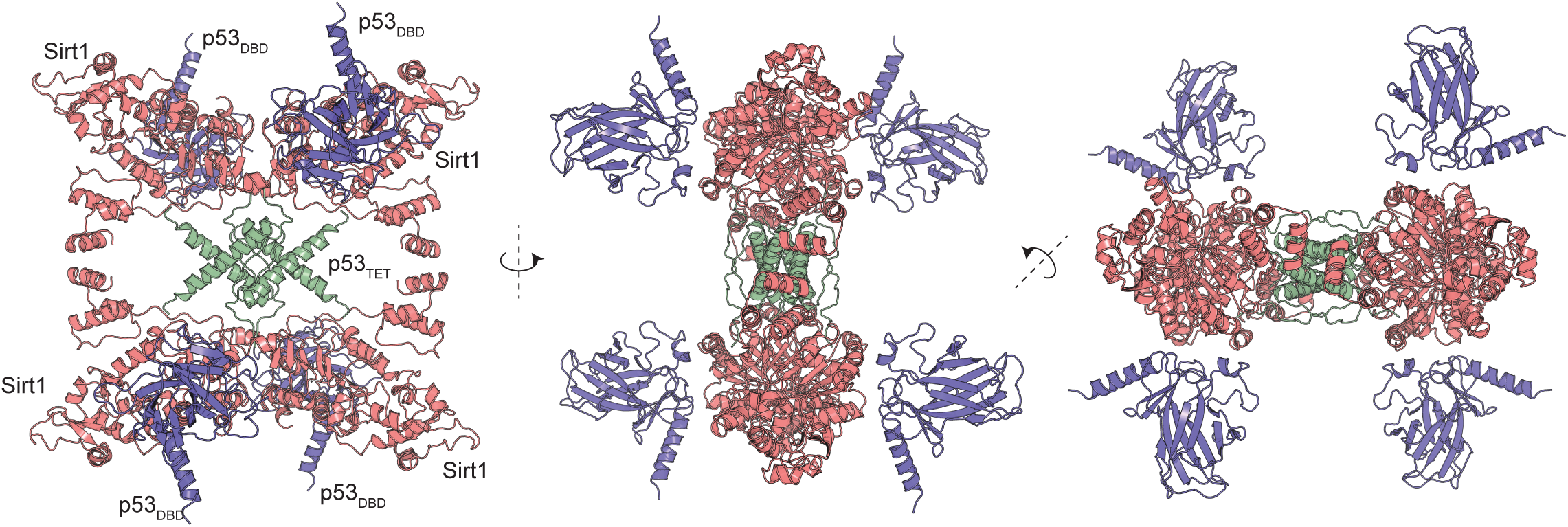

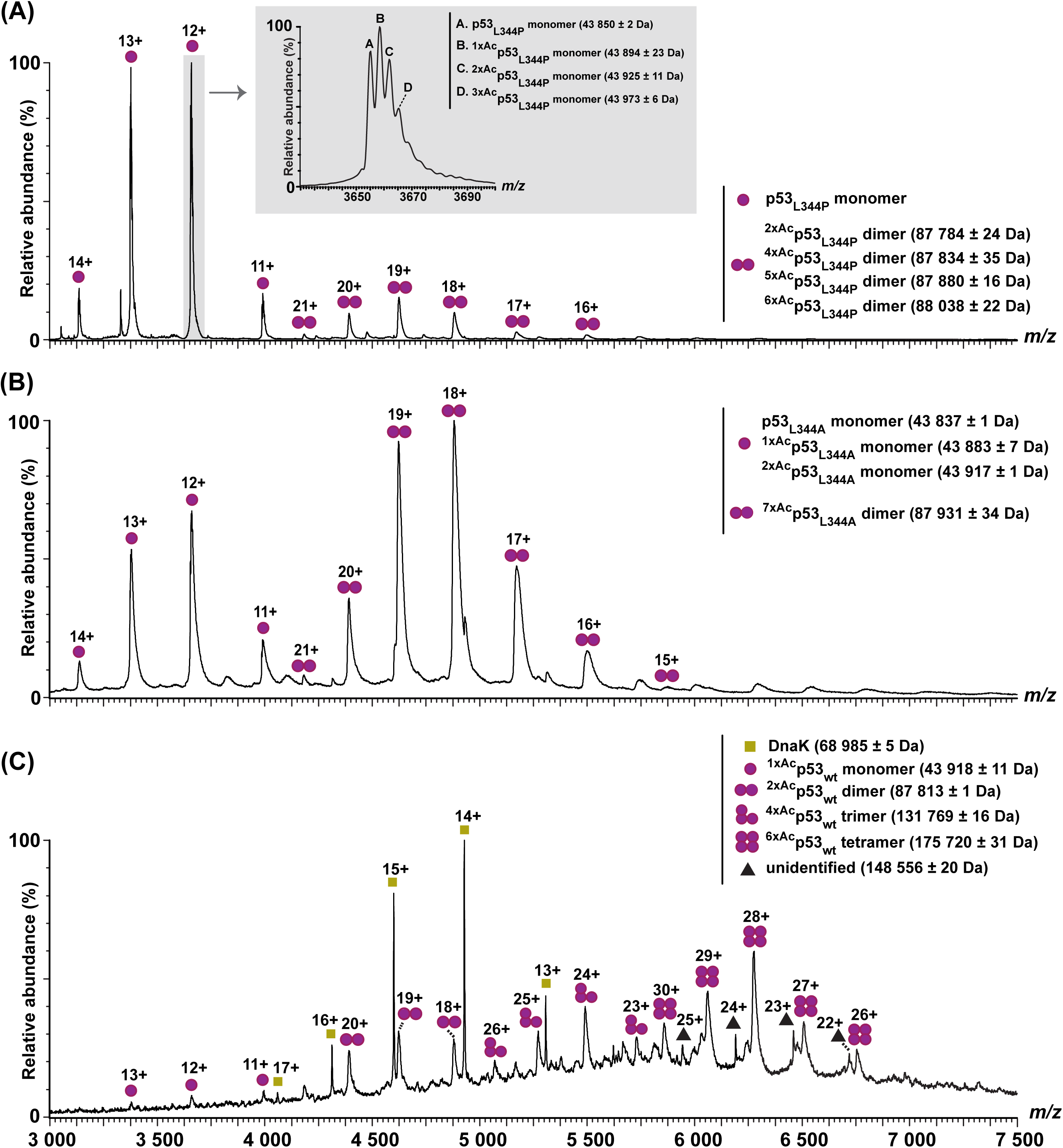

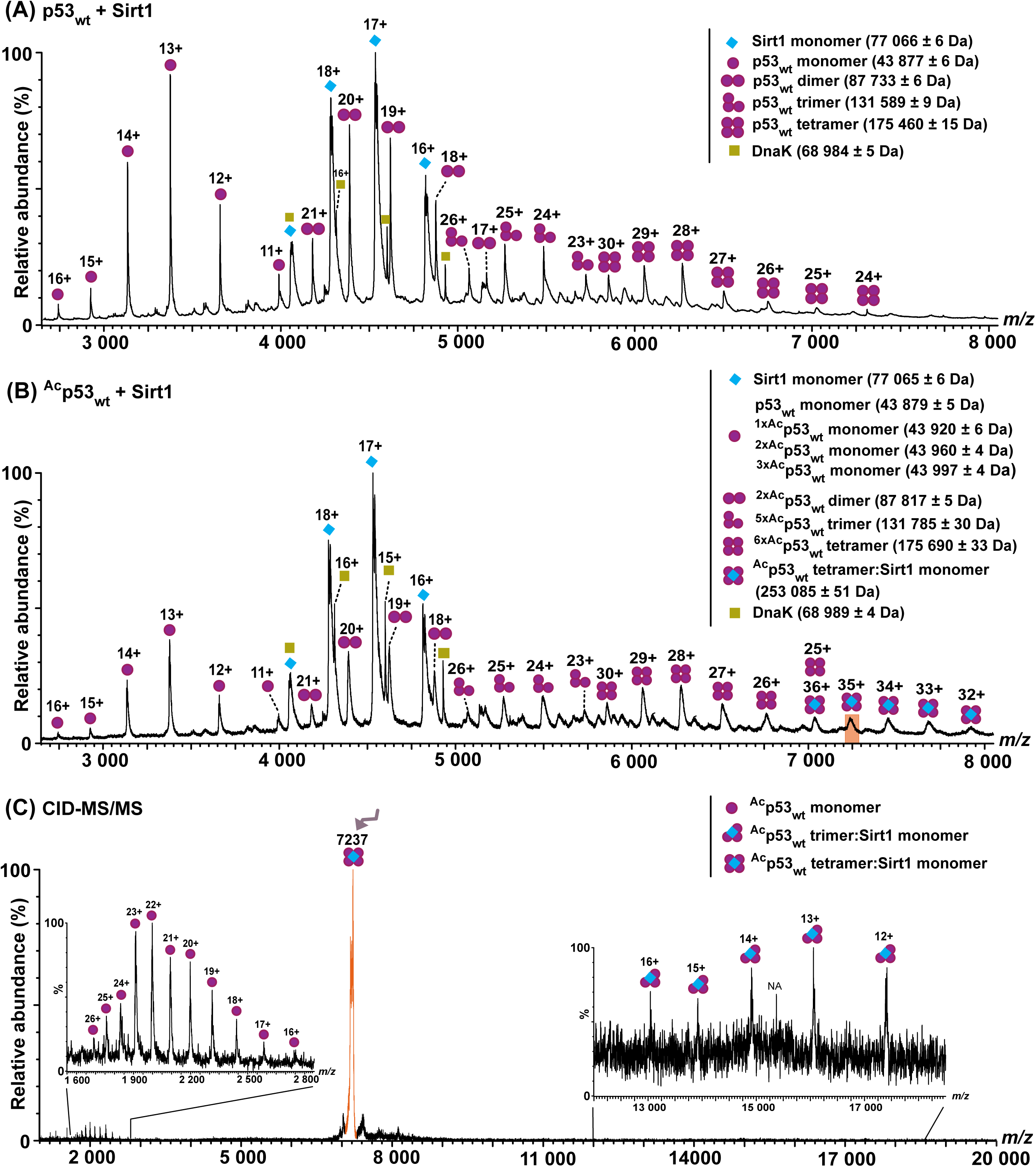

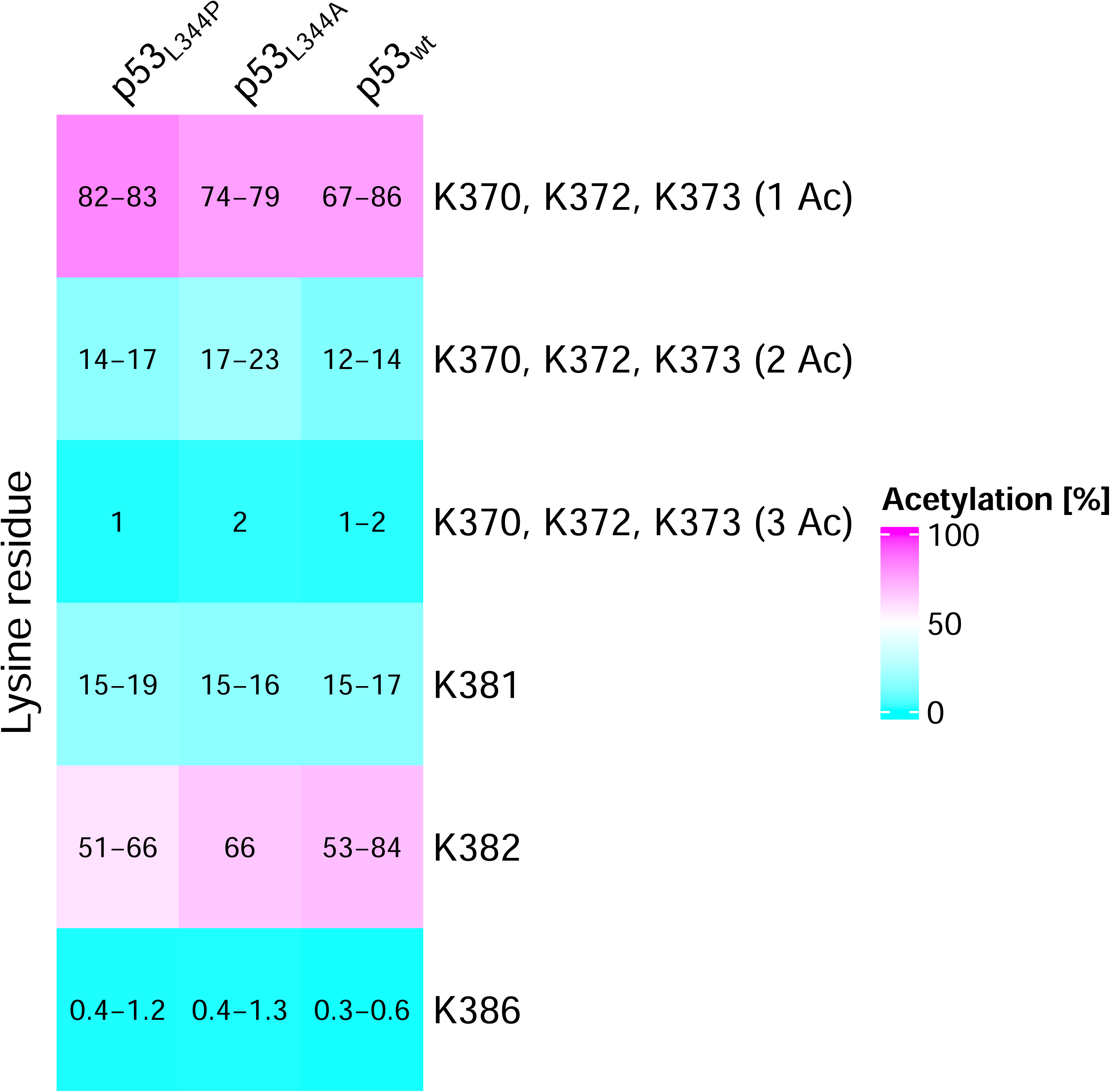

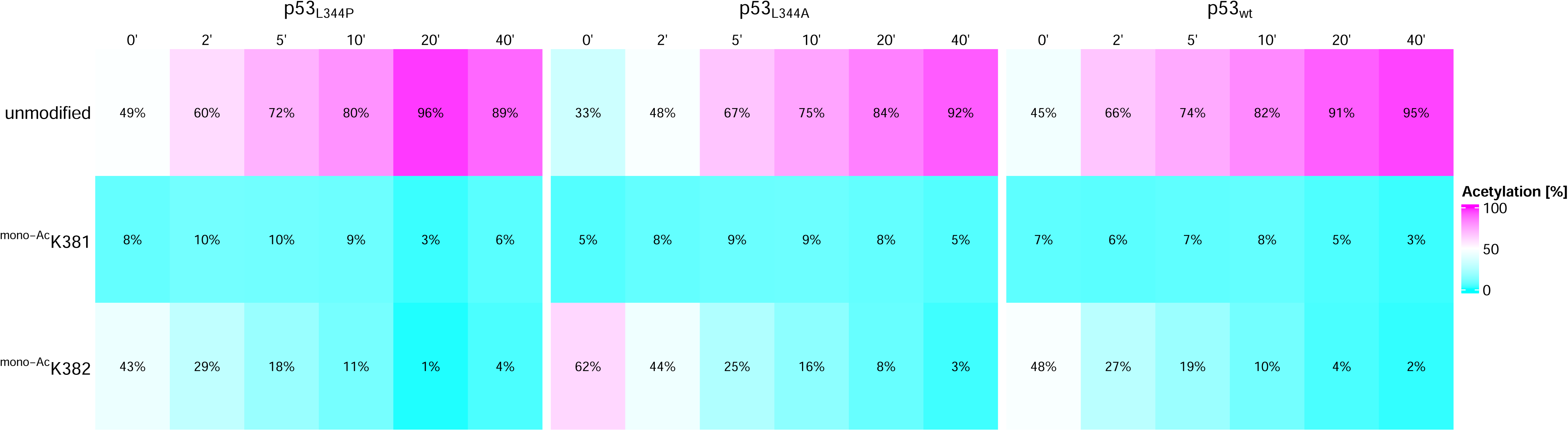

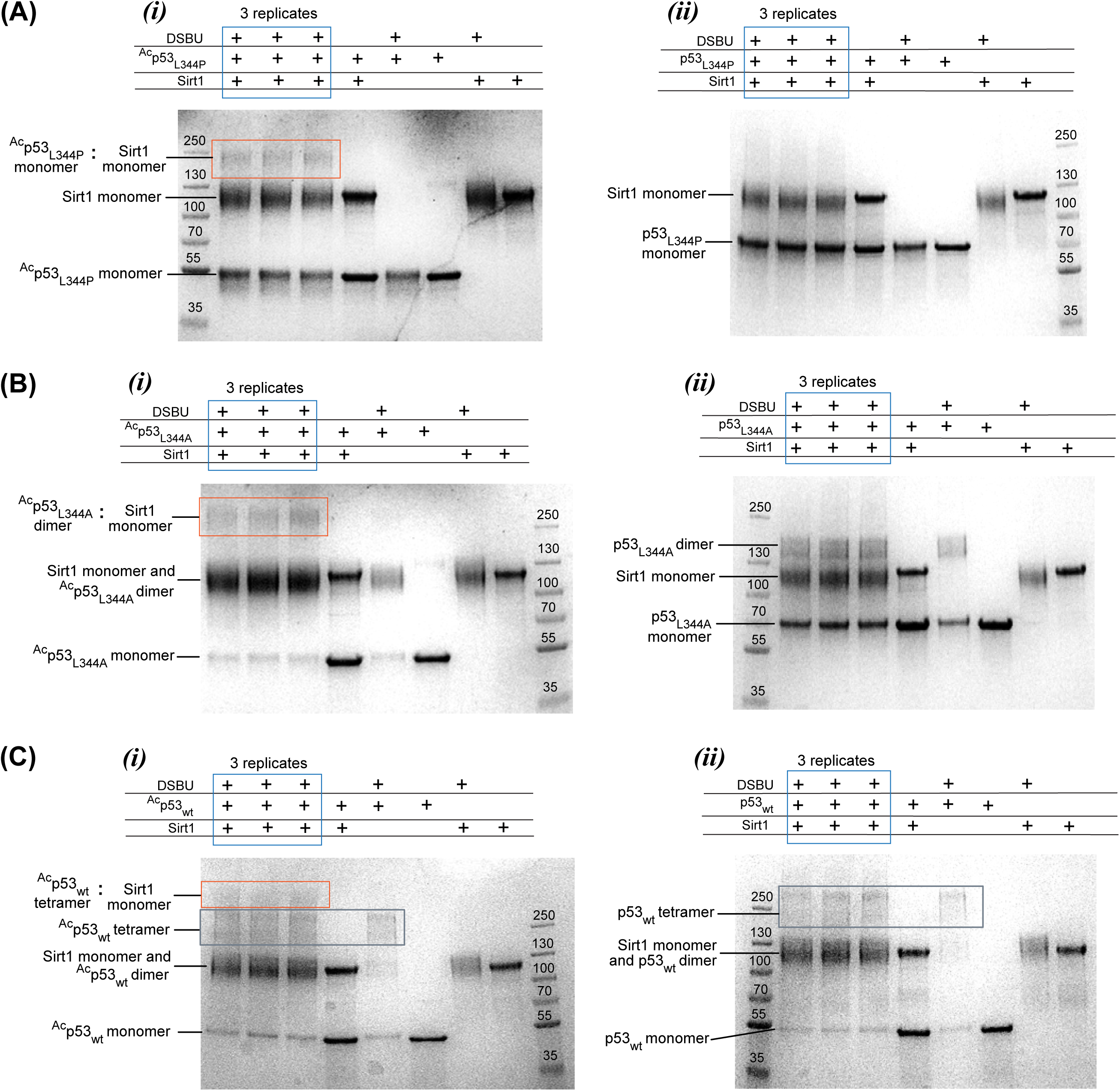

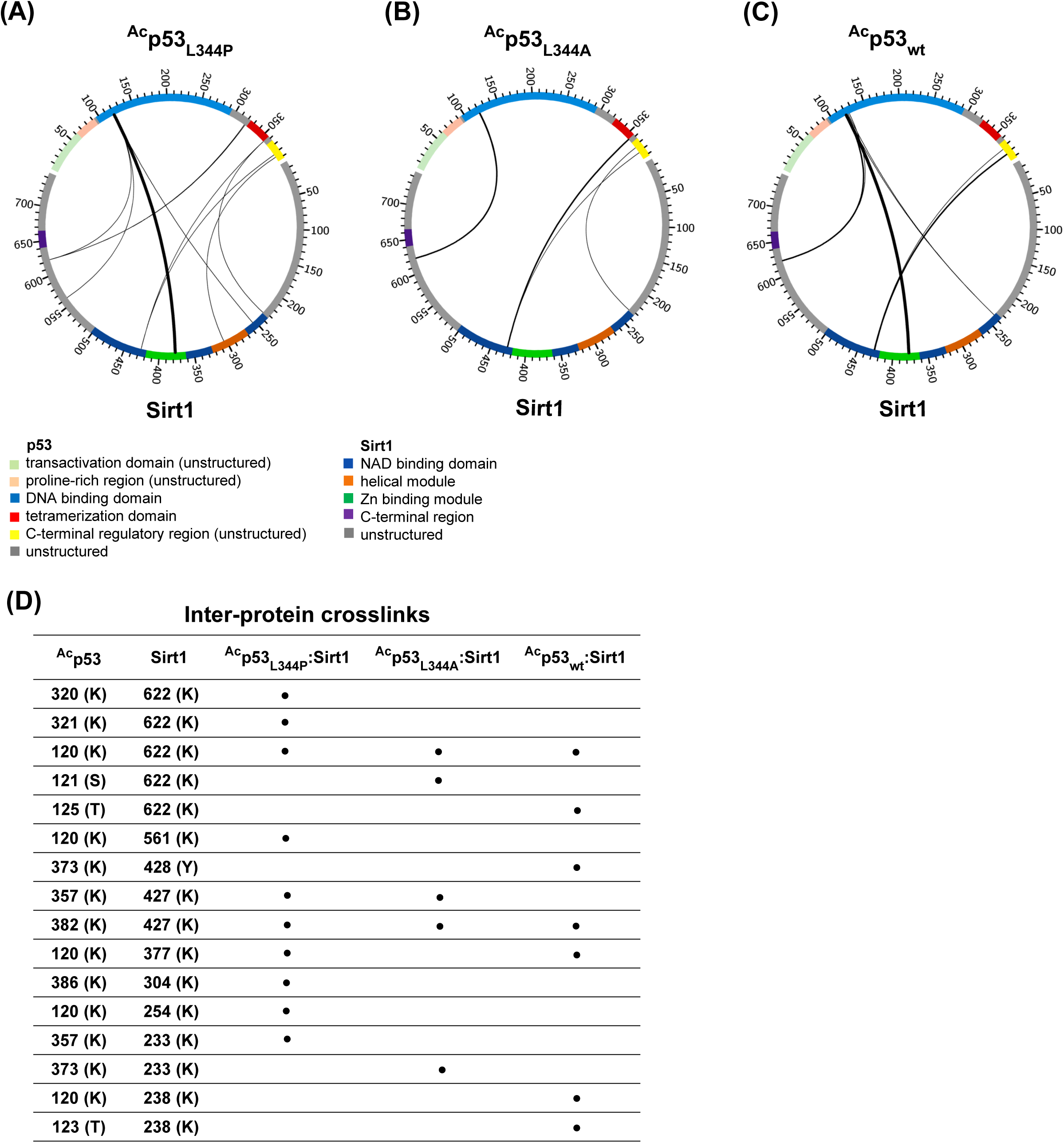

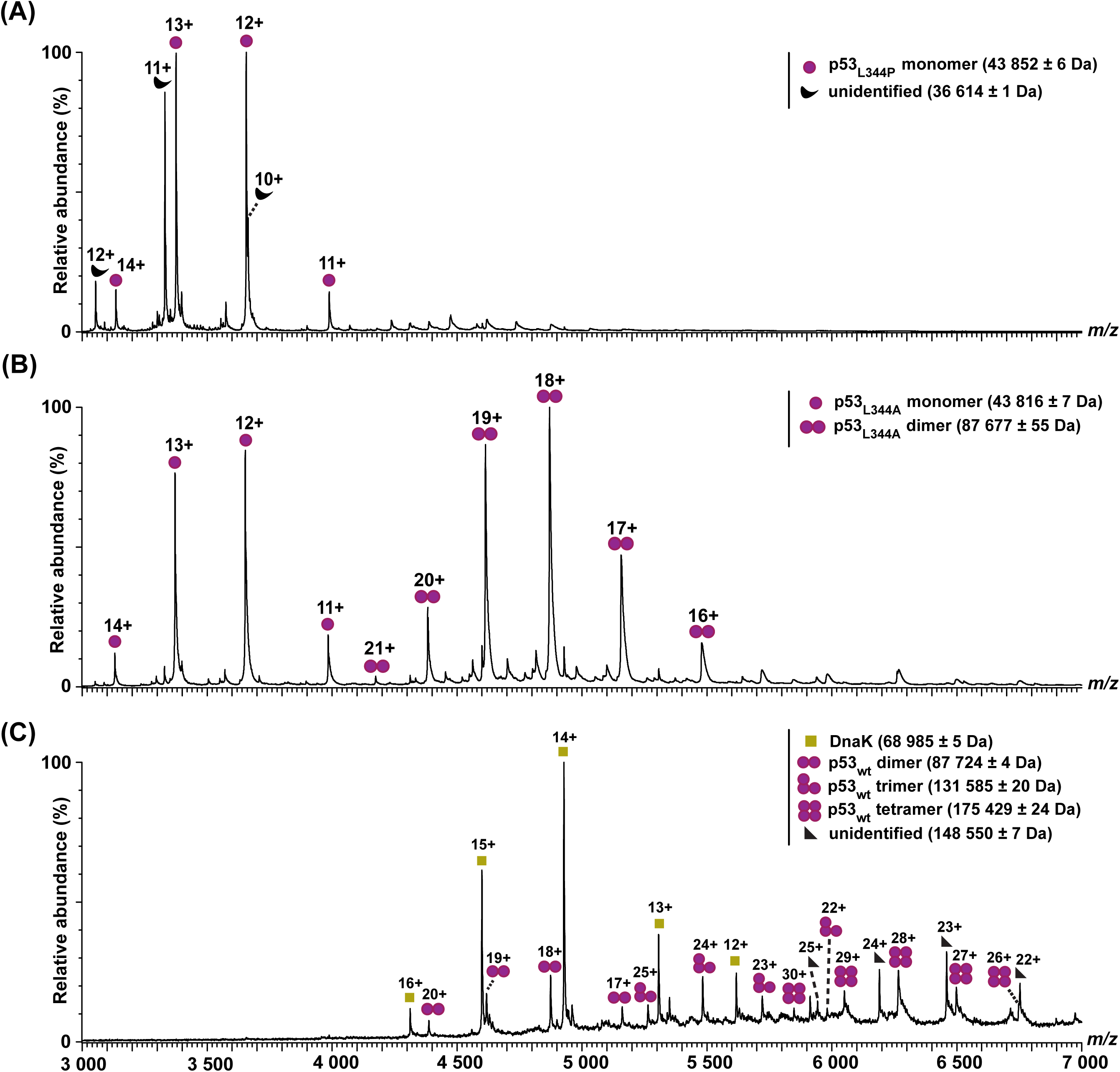

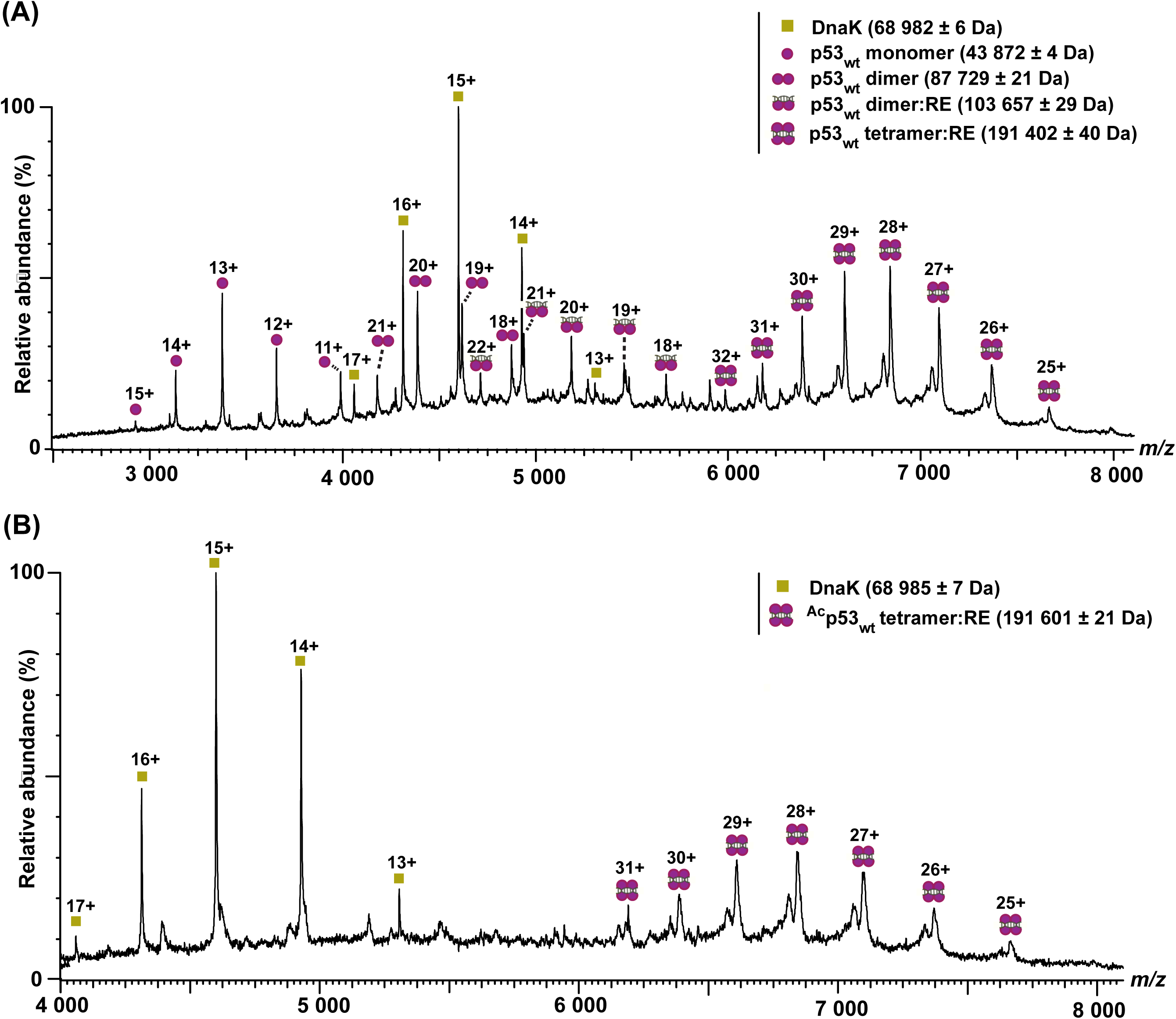

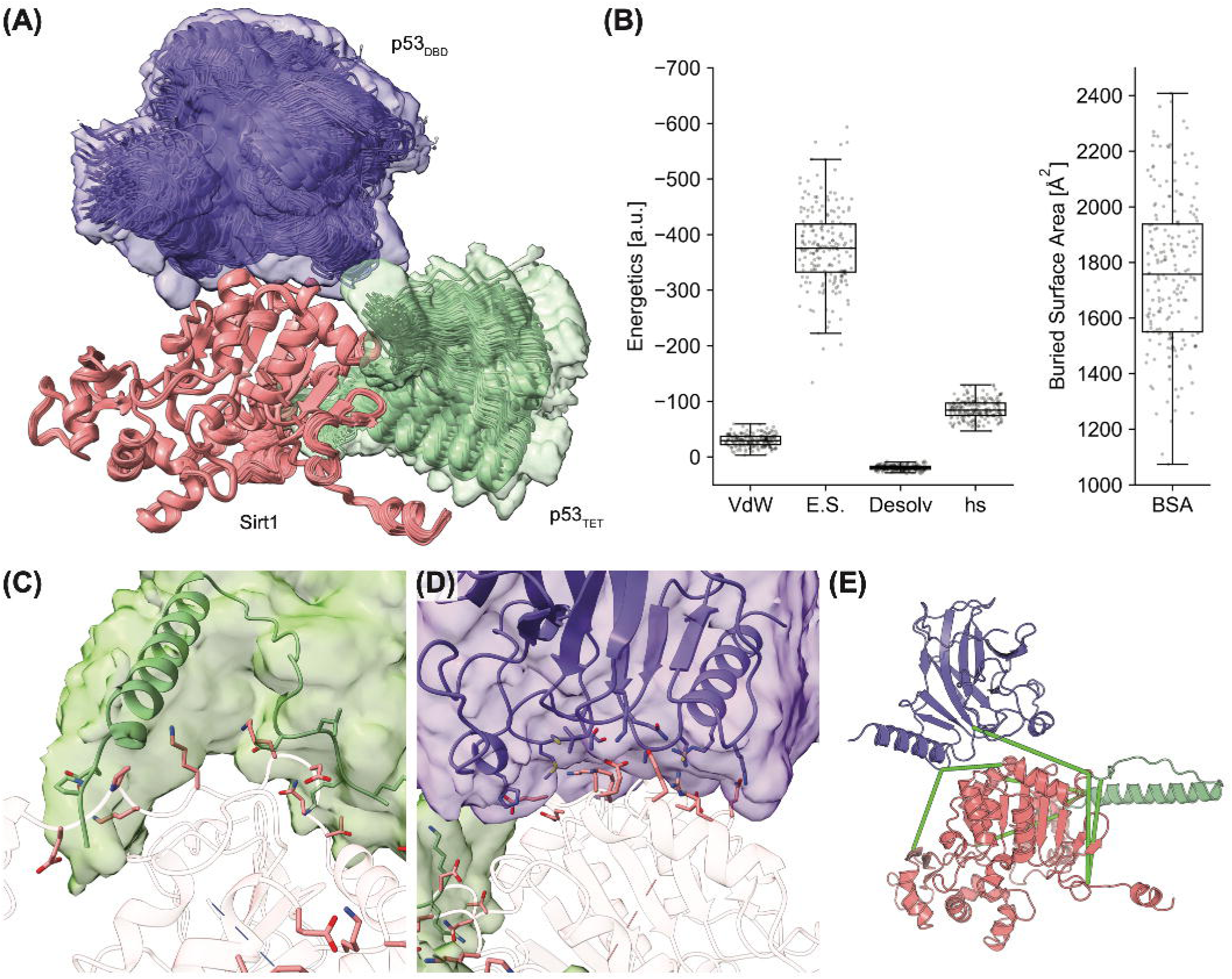

