## Supplementary material for "Revisiting p53:Sirt1 interaction in the light of controlling p53 acetylation levels": Figure Legends

**Figure 1:** Native mass spectra of acetylated full-length human p53 expressed in E. coli. Each p53 monomer is shown as a purple circle, DnaK is shown as yellow square. (A) Acp53L344P forms monomers (charge states 11+ to 14+) with zero, one, two, and three acetylations. Also, dimers are formed with two, four, five, and six acetylations (charge states 16+ to 21+). (B) Acp53L344A forms monomers (charge states 11+ to 14+) with zero, one, and two acetylations as well as dimers (charge states 15+ to 21+) with seven acetylations. (C) Acp53wt forms monomers (charge states 11+ to 13+) with one acetylation, dimers (charge states 18+ to 20+) with two acetylations, trimers (charge states 23+ to 26+) with four, and tetramers (charge states 26+ to 30+) with six acetylations.

**Figure 2:** Native MS confirms acetylation-dependent complex formation between p53_wt_ and Sirt1. (A) For non-acetylated p53_wt_, Sirt1 monomer (charge states 16+ to 19+) as well as p53 monomer (charge states 11+ to 16+), dimer (charge states 17+ to 21+), trimer (charge states 23+ to 26+), and tetramer (charge states 24+ to 30+) are observed. (B) For ^Ac^p53_wt_, Sirt1 monomer (charge states 16+ to 19+) as well as p53 monomer (charge states 11+ to 16+), dimer (charge states 18+ to 21+), trimer (charge states 23+ to 26+), tetramer (charge states 25+ to 30+) are observed. Additionally, the ^Ac^p53_wt_:Sirt1 complex was detected (charge states 32+ to 36+). (C) Collisional activation (CID-MS/MS) of the 35+ charge state of this complex (orange box in B) resulted in the ejection of a ^Ac^p53_wt_ monomer (charge states 16+ to 26+, m/z ~1 600 to ~2 800) from the complex. The remaining ^Ac^p53_wt_:Sirt1 (3:1) complex (charge states 12+ to 16+, m/z ~13 000 to ~17 500) was observed as well.

**Figure 3:** Acetylation profiles of ^Ac^p53 variants co-expressed with p300. Heatmaps depict site-specific acetylation levels (in %) for monomeric p53_L344P_, dimeric p53_L344A_, and tetrameric p53_wt_. Acetylation was detected at lysine residues K370, K372, K373, K381, K382, and K386. Because single acetylation events at K370, K372, and K373 could not be unambiguously assigned to an individual residue, these sites were analyzed collectively, and mono-, di-, and tri-acetylated K370/K372/K373 species are shown as combined states. Ac: acetylation.

**Figure 4:** Time-resolved deacetylation kinetics of ^Ac^p53 variants by Sirt1. The heatmap shows site-specific acetylation levels (in %) of monomeric p53_L344P_, dimeric p53_L344A_, and tetrameric p53_wt_ at lysine residues K381 and K382 (0 to 40 min) upon Sirt1 addition. A gradual loss of mono-acetylated K381 and K382 species is consistent with the accumulation of unmodified p53, indicating progressive deacetylation by Sirt1.

**Figure 5:** SDS-PAGE of p53:Sirt1 cross-linking reaction mixtures. (A) Monomeric p53_L344P_, (B) dimeric p53_L344A_, and (C) tetrameric p53_wt_ were cross-linked with Sirt1 in (i) acetylated or (ii) non-acetylated forms. Cross-linked p53:Sirt1 complexes (orange boxes) were detected only for ^Ac^p53. The apparent molecular weights indicate a 1:1 stoichiometries between ^Ac^p53 and Sirt1, independent of ^Ac^p53’s oligomeric state.

**Figure 6:** Mapping ^Ac^p53:Sirt1 interaction sites by XL-MS. Circos plots illustrate inter-protein cross-linking sites between (A) ^Ac^p53_L344P_ (B) ^Ac^p53_L344A_ and (C) ^Ac^p53_wt_ and Sirt1. Thickness of lines indicate whether a cross-link was found in one, two, or three independent experiments. (D) Summary of cross-linking sites between ^Ac^p53 and Sirt1.

**Figure 7:** Molecular docking of the p53:Sirt1 (1:1) complex. (A) Water-refined ensemble of the molecular docking, aligned to Sirt1 (salmon). All p53_DBD_ (purple) and the p53_TET_ (green) models are shown as cartoon structures. The conformational space is illustrated as surface (n = 188). (B) Energetic distribution of all models, displayed in (A) (n = 188) (Vdw: van der Waals interactions; E.S.: electrostatic score; Desolv: desolvation energy; hs: haddock score; BSA: buried surface area). (C-D) Zoom-in into the protein-protein interaction interface between p53 and Sirt1. Residues in the interface are highlighted as licorice. (C) p53_TET_:Sirt1 interface. (D) p53_DBD_:Sirt1 interface. (E) Cross-link satisfaction, illustrated in the best scored model (based on HADDOCK score). All Cα-Cα distances of cross-links (shown as light green lines) are in the expected distance range (< 30 Å).

**Figure S1:** Native mass spectra of purified non-acetylated p53 variants. (A) p53_L344P_. forms monomers (charge states 11+ to 14+). (B) p53_L344A_ forms monomers (charge states 11+ to 14+) and dimers (charge states 16+ to 21+). (C) p53_wt_. forms dimers (charge states 17+ to 20+), trimers (charge states 23+ to 25+), and tetramers (charge states 26+ to 30+).

**Figure S2:** Native mass spectra of DNA response element (RE)-bound p53_wt_. (A) Non-acetylated p53_wt_ forms monomers (charge states 11+ to 15+) and dimers (charge states 18+ to 21+). In addtion, DNA RE-bound p53_wt_ dimers (charge states 18+ to 22+) and tetramers (charge states 25+ to 32+) were observed. (B) ^Ac^p53_wt_ forms DNA RE-bound tetramer (charge states 25+ to 31+).

**Figure S3:** SDS-PAGE and native MS of Sirt1. (A) SDS-PAGE of samples collected at different stages of Sirt1 purification. (B) Native mass spectrum depicting purified intact Sirt1 (blue diamonds) in monomeric (charge states 15+ to 19+) and dimeric (charge states 23+ to 28+) state.

**Figure S4:** Native MS confirms acetylation-dependent complex formation between p53_L344P_ and Sirt1. (A) For non-acetylated p53_L344P_, Sirt1 monomer (charge states 15+ to 20+) and dimer (charge states 23+ to 28+) as well as p53_L344P_ monomer (charge states 12+ to 15+) was observed. (B) For ^Ac^p53_L344P_, Sirt1 monomer (charge states 16+ to 20+) and dimer (charge states 24+ to 28+), p53 monomer (charge states 12+ to 15+), and the ^Ac^p53_L344P_:Sirt1 complex (1:1) (charge states 21+ to 24+) have been detected. (C) Collisional activation (CID-MS/MS) of the 23+ charge state of this complex (orange box in B) resulted in the ejection of a ^Ac^p53_L344P_ monomer (charge states 12+ to 17+, m/z ~2 600 to ~3 700) from the complex. The remaining Sirt1 (charge states 7+ to 9+, m/z ~7 500 to ~11 000) was observed as well.

**Figure S5:** Native MS confirms acetylation-dependent complex formation between p53_L344A_ and Sirt1. (A) For non-acetylated p53_L344A_, Sirt1 monomer (charge states 14+ to 19+) as well as p53_L344A_ monomer (charge states 11+ to 14+) and dimer (charge states 16+ to 20+) were observed. (B) For ^Ac^p53_L344A_, Sirt1 monomer (charge states 14+ to 18+) and dimer (charge states 23+ to 26+), p53 monomer (charge states 11+ to 14+) and dimer (charge states 17+ to 20+), as well as the ^Ac^p53_L344A_:Sirt1 complex (2:1) (charge states 24+ to 27+) have been detected. (C) Collisional activation (CID-MS/MS) of the 27+ charge state of this complex (orange box in B) resulted in the ejection of a ^Ac^p53_L344A_ monomer (charge states 15+ to 20+, m/z ~2 200 to ~3 000) from the complex. The remaining ^Ac^p53_L344A_:Sirt1 (1:1) complex (charge states 9+ to 12+, m/z ~10 000 to ~13 500) was observed as well.

**Figure S6:** AlphaFold2 model preparation for molecular docking. Atomic models are colored according to the plDDT confidence score (see color bar). Both initial models were retrieved from AlphaFold-DB. Low-confidence regions outside of folded domains were removed to allow for rigid-body docking. (A) p53. From the AlphaFold2 model, the p53_DBD_ consists of residues 95–295, p53_TET_ comprises residues 320–360. (B) Sirt1. The core region of Sirt1 (residues 183–670), with residues 501–633 missing, were selected and used for rigid-body docking.

**Figure S7:** Superimposition of the docked p53*:*Sirt1 complex on the p53_TET_ crystal structure. Four 1:1 complexes of p53*:*Sirt1 were aligned on the published p53_TET_ structure (PDB 1c26). There is no clash between the ordered domains, but the p53_DBD_ structures are disconnected by 84-114 Å. The first rotation is 90° at the y-axis, the second rotation 90° at the z-axis (green: p53_TET_; purple: p53_DBD_; salmon: Sirt1).
